# Quantification of macromolecular biomass composition for constraint-based metabolic modeling

**DOI:** 10.1101/2021.08.20.457062

**Authors:** Vetle Simensen, Christian Schulz, Emil Karlsen, Signe Bråtelund, Idun Burgos, Lilja Brekke Thorfinnsdottir, Laura García-Calvo, Per Bruheim, Eivind Almaas

## Abstract

Genome-scale metabolic models (GEMs) are mathematical representations of metabolism that allow for *in silico* simulation of metabolic phenotypes and capabilities. A prerequisite for these predictions is an accurate representation of the biomolecular composition of the cell necessary for replication and growth, implemented in GEMs as the so-called biomass objective function (BOF). The BOF contains the metabolic precursors required for synthesis of the cellular macro- and micromolecular constituents (e.g. protein, RNA, DNA), and its composition is highly dependent on the particular organism, strain, and growth condition. Despite its critical role, the BOF is rarely constructed using specific measurements of the modeled organism, drawing the validity of this approach into question. Thus, there is a need to establish robust and reliable protocols for experimental condition-specific biomass determination. Here, we address this challenge by presenting a general pipeline for biomass quantification, evaluating its performance on *Escherichia coli* K-12 MG1655 sampled during balanced exponential growth under controlled conditions in a batch-fermentor set-up. We significantly improve both the coverage and molecular resolution compared to previously published workflows, quantifying 91.6% of the biomass. Our measurements display great correspondence with previously reported measurements, and we were also able to detect subtle characteristics specific to the particular *E. coli* strain. Using the modified *E. coli* GEM *i*ML1515a, we compare the feasible flux ranges of our experimentally determined BOF with the original BOF, finding that the changes in BOF coefficients considerably affect the attainable fluxes at the genome-scale.

## Introduction

The increasing availability of large-scale omics data has propelled the study of complex biological systems, pushing the field of systems biology to the forefront of cutting-edge biological research [1, 2]. Central to this development is the realization that biology is best understood not merely by considering its individual constituents, but rather investigating the emergent properties of the system as a whole. One of the predominant subfields of systems biology is the *in silico* study of metabolism using genome-scale metabolic models (GEMs) [3–6]. Here, the genetically encoded metabolic potential of an organism is used to construct a stoichiometric network of biochemical transformations. The steady-state fluxes of the metabolic system can subsequently be calculated using approaches such as flux balance analysis [7]. These flux phenotypes are usually computed by maximizing growth, assuming optimal biomass production to be a reasonable cellular objective [8].

Growth in these models is implemented as a pseudo-reaction called the biomass objective function (BOF), whose reactants are the metabolic precursors needed to generate the molecular constituents of the cell. By appropriately scaling the stoichiometric coefficients of these precursors using experimental biomass measurements, the flux through the BOF directly corresponds to the specific growth rate, allowing for quantitative predictions of growth phenotypes [9, 10]. While experimental data on the condition-dependent biomass compositions of some well-studied organisms are available [11, 12], this is commonly lacking for most organisms. The usual strategy has therefore been to either adopt existing organism-specific biomass compositions from different conditions or employ parts or the whole composition from another organism entirely [13, 14]. This, however, is a sub-optimal approach as the biomass composition depends on the particular organism and strain [15, 16]. The biomass composition is also not static, but is rather continually adjusted in response to changing environmental conditions [12].

In many instances, the predicted flux phenotypes of these models have been shown to be highly susceptible to variations in the biomass composition [17]. Dikicioglu *et al*. [18] demonstrated how the predicted flux distributions of a *Saccharomyces cerevisiae* GEM were sensitive to changes in the stoichiometric coefficients of the BOF within experimentally determined bounds. Lakshmanan *et al*. [17] observed a similar sensitivity when varying the biomass composition in GEMs of eight different yeast species, showcasing the impact on both growth rate and gene essentiality predictions to alterations in the biomass constituents and stoichiometries. High-quality, condition-dependent biomass measurements are therefore necessary in order to enable accurate phenotypic predictions using a constraint-based modeling framework. This realization has sparked multiple initiatives for the measurement of biomass compositions [15, 16, 19, 20], and ways to address the challenge of integrating variable biomass compositions into GEMs [21].

The macromolecular composition of an organism can be quantified using a range of experimental procedures. The DNA, RNA, and carbohydrate contents are usually measured by spectroscopic methods, while the total cellular protein content is quantified by acid hydrolysis followed by high-performance liquid chromatography (HPLC) [15]. Total lipid content is commonly obtained by extraction and gravimetric quantification, whereas the lipid class and fatty acid compositions are measured using various mass spectrometry-based (MS) approaches [22]. Recently, Beck *et al*. [15] conducted a literature review and developed a step-by-step protocol for quantifying each macromolecular biomass component, evaluating its applicability on multiple bacterial samples. Although the pipeline exhibited comparable efficiencies in between the bacterial species, the total macromolecular contents covered by the methods were approximately 65%, necessitating extensive loss-adjustment by normalization in order to construct a BOF. The quantification of the biochemically diverse carbohydrates was also quite limited in molecular resolution, only measuring the total biomass contents. Another workflow proposed by Long and Antoniewicz [19] applied gas chromatography/mass spectrometry (GC/MS) as a single analytical platform to absolutely quantify the macromolecular composition of amino acids, RNA, fatty acids, and glycogen in *Escherichia coli*. While they obtained an impressive overall coverage of the total biomass, the quantification was based on isotope ratio analysis, requiring the cells to be fully labeled with ^13^C as well as a supplementation of multiple standard compounds for quantification.

Here, we present a concise pipeline for accurate high-coverage absolute biomass quantification, significantly increasing both the yield and molecular resolution in comparison to previous work. *E. coli* K-12 MG1655 was grown aerobically in a defined glucose minimal medium using a batch fermentor setup. We evaluated the performance of our pipeline, obtaining high consistency with previously reported values. Specifically, we extend and adjust the workflow of Beck *et al*. [15] by improving the resolution of the carbohydrate analysis using liquid chromatography UV and electrospray ionization ion trap technique (HPLC-UV-ESI-MS/MS) [23]. Furthermore, we monitored fermentation parameters to ensure stable and controlled conditions throughout the experiment, allowing for biomass sampling in the exponential growth phase. With these enhancements, we obtain an overall mass coverage of 91.6%, proving the pipeline to be an important milestone towards absolute biomass quantification for computational biology applications.

To explore the modelling impact of the BOF generated from our experiments, we used flux variability analysis (FVA) on the *i*ML1515 GEM [24]. Specifically we assessed how, in an aerobic minimal glucose medium, the experimentally determined BOF (hereafter *eBOF*) differed in predicting feasible flux ranges compared to the BOF included in the *i*ML1515 GEM (hereafter *mBOF*). The eBOF is made in a modified version of the GEM named *i*ML1515a. We find that even though both the mBOF and the eBOF is supposed to originate from the same experimental conditions, they differ considerably in their coefficient values as well as in their phenotypic predictions of genome-scale flux ranges.

## Materials and methods

### Strain, media and culture conditions

We prepared *E. coli* strain K-12 MG1655 (700926™, ATCC^®^) glycerol stock solutions by growing the organism on an LB agar plate and selecting a single colony. The LB agar contained the following: 10 g L^−1^ peptone, 5 g L^−1^ yeast extract, 5 g L^−1^ NaCl, and 15 g L^−1^ agar. An overnight culture in minimal M9 glucose medium was then aliquoted with 25% glycerol and stored at −80 °C. To prepare the inoculum for the fermentation, an aliquote of the glycerol stock solution was grown overnight in an incubator with shaking (37 °C, 200 rpm) using 100 mL of a standard M9 minimal salts medium with glucose as the sole carbon source in a 500 mL baffled shake flask. The minimal M9 medium had the following composition: 0.4% (w/v) glucose, 1 mM MgSO_4_, 18.7 mM NH_4_Cl, 8.5 mM NaCl, 22.0 mM KH_2_PO_4_, 33.7 mM Na_2_HPO_4_, and 0.2% (v/v) trace mineral solution. The concentration of trace minerals in the finished medium was 35.9 mM FeSO_4_ · 7 H_2_O, 4 mM CuSO_4_ · 5 H_2_O, 7.8 mM ZnSO_4_ · 7 H_2_O, 1.9 mM MnCl_2_ · 4 H_2_O, 0.08 mM (NH_4_)_6_Mo_7_O_24_ · 4 H_2_O, 0.2 mM CoCl_2_ · 6 H_2_O, and 13.6 mM CaCl_2_ 2 H_2_O dissolved in 1 M HCl. The cells were cultured in a 3 L

Eppendorf NewBrunswik BioFlo 115 bioreactor in batch setup, using 1 L of a modified minimal M9 medium containing 1% (w/v) glucose, 3 mM MgSO_4_, 93.5 mM NH_4_Cl, 8.5 mM NaCl, 11.5 mM KH_2_PO_4_, and 0.2% (v/v) of the same trace mineral solution used in the preculture. The pH electrode was calibrated using a two-step calibration with pH 4 and pH 7 pre-mixed solutions. The dissolved oxygen (DO) electrode was calibrated to 0% by flushing the electrode for 10 min with nitrogen gas, and to 100% dissolved oxygen at 37 °C in the fermentor medium after 30 min with ≈ 500 mL min^−1^ air inflow at 500 rpm stirring. Stirring was then coupled to DO, ensuring DO≥ 40%.

The off-gas was analyzed with an Eppendorf DASGIP GA4 gas analyzer, allowing for continuous monitoring of the O_2_ consumption and CO_2_ production. The gas in- and outflow from the fermentor was sterile filtered by employing 0.2 µm filters. The pH was kept constant at pH 7 using 4 M NaOH by automatic titration. Foam production was controlled by the manual addition of silicone-polymer based Antifoam. The bioreactor was inoculated with 1% inoculum. The cells were harvested during exponential growth, centrifuged for 5 min (3645 g, 4 °C), and washed two times in 0.9% NaCl solution, followed by one washing step with MQ water. The pellets were frozen at −80 °C and lyophilized for 3 d. The resulting cell dry mass was then used for the respective protocols.

### Medium analysis

The quantification of medium constituent concentrations was performed by NMR, using ERETIC2 [25] in the Bruker TopSpin 4.0.8 software. The protocol is based on Søgaard *et al*. [26]. At specific time points, medium was collected and sterile filtered, and 2.5 mL were stored at −20 °C before lyophilization and re-hydration in 600 µL D_2_O-TSP (0.75%) solution. 500 µL were transferred into a 5 mm NMR tube and analyzed in a 400 MHz (14.7T) Bruker NMR spectrometer applying the D_2_O solvent setting (^1^H NMR, noesyggpr1d). The acquisition parameters were 4 dummy scans, 32 scans, SW 21.0368 ppm, O1 1880.61 Hz, TD 65536, TE 300.0 K, D1 4 s, AQ 3.892 838 5 s, and P1 was calibrated for each sample to ensure accurate quantification. A 70 mM creatine solution (in D_2_O) was used as an external standard, utilizing the singlet at ∼ 3 ppm for quantification. As an example, for the glucose quantification the *α*-Glucose doublet at ∼ 5.2 ppm was used, which accounts for 36% of the glucose [27]. The peaks of formate, acetate, succinate, and lactate were identified based on the reference ^1^H-NMR spectra available in the Human Metabolome Database (HMDB) [28–30] and the software program Chenomx [31], as well as literature by Fan [32].

### Protein

The total cellular protein content was measured by acid hydrolysis, followed by amino acid derivatization and quantification by HPLC based on the protocol described by Noble *et al*. [33]. Aliquots of ∼ 10 mg lyophilized cell dry mass were re-hydrated in 5 mL 6 M hydrochloric acid in a 25 mL Schott flask, boiled for 24 h at 105 °C, and allowed to cool to handling temperature before neutralizing with 5 mL 6 M NaOH. The samples were again cooled to handling temperature before sterile filtering. Using different dilutions (see S3 File) of the filtered samples, the amino acids were quantified by reverse-phase HPLC analysis. The samples and standards were derivatized with OPA (o-Phthaldialdehyde Reagent Solution). We used a Waters Nova-Pak C18 4 µm column (3.9×150 mm) with an RF2000 detector set to 330 nm excitation and 438 nm emission wavelength. Further, we employed two mobile phases (phase A: methanol and phase B: 0.08 M CH_3_COONa adjusted to pH 5.90 with concentrated CH_3_COOH and 2% tetrahydrofuran just before usage) with a flow rate of 0.9 mL min^−1^ for all gradients, the injection volume was 20 µL. The gradients in 29 min run time were [0 to 15) min 0.25 A and 0.75 B, [15 to 22) min 0.5 A and 0.5 B, [22 to 26) min 0.8 A and 0.2 B, [26 to 27) min 1 A and 0 B, and [27 to 29) min 0.25 A and 0.75 B. Not all amino acids were directly measured, either due to partial or complete degradation during hydrolysis (e.g. methionine, cysteine). The levels of these amino acids were therefore estimated based on a linear regression of measured amino acid mass fractions and their corresponding prevalence in protein-coding genes. The amino acids with overlapping retention times were treated in the same fashion (i.e. glycine and arginine), as were the levels of glutamine and asparagine which are deamidated to glutamate and aspartate, respectively, during acid hydrolysis [34].

### RNA

The cellular proportion of RNA was quantified spectrophotometrically using the protocol described by Benthin *et al*. [35]. ∼30 mg of lyophilized biomass were washed three times with 3 mL 0.7 M HClO_4_ by vortexing and centrifuging at 2880 g for 10 min at 4 °C, discarding the supernatant between washes. The resulting cell pellet was re-suspended in 3 mL 3 M KOH and incubated in a water bath at 37 °C for 1 h, shaking at 15 min intervals. The samples were cooled and 1 mL 3 M HClO_4_ was added before centrifuging at 2880 g for 10 min at 4 °C, decanting the supernatant into a 50 mL polypropylene centrifuge tube. The pellet was washed (re-suspended and centrifuged) twice with 4 mL 0.5 M HClO_4_, before the supernatant was decanted into the 50 mL tube. 3 mL 0.5 M HClO_4_ was added to the collected sample and centrifuged to remove any precipitates of KClO_4_. The RNA concentration was measured via UV-visible spectroscopy against the reference solvent using the NanoDrop [36]. The levels of the individual ribonucleotides were estimated based on the monomeric composition of rRNA-encoding genes in *E. coli* strain K-12 MG1655 (GenBank accession number U00096.3 [37]), as these constitute approximately 81% of the total RNA content in *E. coli* [38].

### DNA

DNA was extracted using the protocol described in Wright *et al*. [39]. ∼ 10 mg lyophilized biomass were dissolved in 600 µL lysis buffer (9.34 mL TE buffer containing 10 mM Tris-Cl (pH 8.0) and 1 mM EDTA (pH 8.0), 600 µL 10% SDS, and 60 µL proteinase K (20 mg mL^−1^)) and incubated at 55 °C for 30 min before cooling to room temperature. 600 µL phenol/chloroform (1:1 v/v) were added and mixed well. The samples were centrifuged for 5 min at 12 044 g (max speed) in a table centrifuge at room temperature and the upper aqueous phase was transferred to a separate tube. The addition of phenol/chloroform and the subsequent mixing and centrifuging was repeated twice, each round pooling the aqueous phases. An equal volume of chloroform was added to the aqueous phase and the solution was mixed well. The tube was centrifuged for 5 min at max speed in a table centrifuge at room temperature. To precipitate the DNA, the aqueous phase was separated, and mixed gently with 40 µL NaCH_3_COO and 1 mL ice cold ethanol (99%), then incubating at −20 °C for 30 min. The sample was centrifuged for 15 min at max speed in a table centrifuge. The supernatant was discarded and the pellet was rinsed using 1 mL ethanol (70%). The tube was centrifuged for 2 min (table centrifuge, max speed), before carefully discarding the supernatant and air-drying the DNA pellet. The pellet was re-suspended in 50 µL TE buffer and 1 µL RNAase A was added before incubating for 15 min at 37 °C. The concentration of dsDNA was measured by UV-visible spectroscopy using the NanoDrop [36]. The relative distribution of individual deoxyribonucleotides was estimated based on the genome sequence of *E. coli* strain K-12 MG1655 (GenBank accession number U00096.3 [37]).

### Carbohydrate

The total carbohydrate content was measured by order at the Technical University of München in Germany following the protocol described by Rühmann *et al*. [40]. Briefly, cell dry mass (2 mg L^−1^) was hydrolyzed in 4 M trifluoroacetic acid for 90 min at 121 °C and derivatized with 1-phenyl-3-methyl-5-pyrazolone. The carbohydrate analysis was then performed via HPLC-UV-ESI-MS/MS. This assures high-quality identification and quantification of a wide range of chemically diverse carbohydrate monomers and dimers.

### Lipid

Lipids were quantified gravimetrically following a chloroform/methanol extraction protocol [41, 42]. ∼ 40 mg lyophilized cell dry mass were re-hydrated by adding 0.15 mL water and vortexed briefly at low rpm. The re-hydrated cells were homogenized in a homogenizer at 6500 rpm for 20 s intervals, 2 cycles, along with ∼ 0.5 g zirconium beads (1.4 mm) and 0.4 mL methanol. The samples were kept on ice between runs. 0.8 mL chloroform were added before vortexing for 20 min, subsequently adding 0.1 mL water and vortexing again for 10 min. The sample tubes were centrifuged for 4 min at max speed using a table centrifuge, after which the lower chloroform phase was transferred to a separate tube, before repeating the chloroform extraction with 0.6 mL chloroform.

Finally, the chloroform was allowed to completely evaporate (≈ 24 h to 36 h). The total lipid content was quantified by weighing, and corrected using blanks, as well as loss-adjusted for incomplete retrieval during extraction.

### Construction of a novel BOF

Using our experimental measurements, we constructed a novel BOF (eBOF) for the *E. coli* GEM *i*ML1515. For a detailed description of this process, see Supplementary material S2 File and S3 File. Briefly, the stoichiometric coefficients of the existing macromolecular precursors in the *i*ML1515 model BOF, here termed *mBOF*, were adjusted to reflect our measurements and normalized to a molar mass (g mmol^−1^) of unity. In cases where the precursors themselves were complex biomolecules (e.g. LPS (lipopolysaccharides)), their contents were estimated using our measurements and their molecular composition.

### Flux comparison by flux variability analysis

To assess the impact on phenotypic predictions, we performed flux variability analysis (FVA) [43, 44] on *i*ML1515 using both mBOF and eBOF as the cellular objectives, with an optimality constraint of 100%. For the sake of simplicity and preventing bias, the modeling was performed using the default exchange rates provided with the model, except that the lower bound on the exchange reaction for cobalamin (EX cbl1 e) was adjusted from 0 to − 1000 mmol gCDW^−1^, as the mBOF supplied with the model (and consequently the eBOF) would not grow without.

To compare the resulting minimal and maximal reaction fluxes, we calculated the fractional overlap, *ξ*, of the corresponding flux range. For a given reaction flux *v*_*j*_ with minimal and maximal fluxes *α*_*min,j*_ and *α*_*max,j*_ for eBOF and minimal and maximal fluxes *β*_*min,j*_ and *β*_*max,j*_ for mBOF, *ξ* is defined as

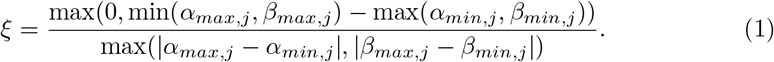

We also calculated the relative value of the eBOF coefficients to those of mBOF, *S*_*r*_, given by

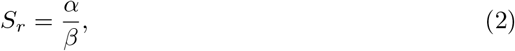

where *α* and *β* are the stoichiometric coefficients (mmol gCDW^−1^) of a given biomass component in eBOF and mBOF, respectively. All simulations were performed in Matlab 2020a [45] using the COBRA toolbox v3 [46] with Cplex [47] as solver.

## Results

### Batch-fermentation and biomass sampling

To obtain the samples for biomass compositional quantification, we cultured *E. coli* K-12 MG1655 aerobically in a bioreactor in batch set-up using a defined glucose minimal medium (see section Materials and methods, subsection Strain, media and culture conditions). The cells exhibited a specific growth rate of 0.71 h^−1^ (generation time 58.7 min), resulting in a cell density of ∼ 2.6 gCDW L^−1^ when sampled during the balanced exponential growth phase at approximately 7.5 h. The corresponding growth curve and time-course fermentation profile is presented in Fig 1 (data from S4 File, full time-course profile can be seen in S4 Fig). *E. coli* displayed a prototypical respiro-fermentative metabolism with extracellular accumulation of the mixed-acid fermentation products acetate, lactate, formate, and succinate. Following the sampling, washing and subsequent lyophilization, the dry cell mass was analyzed using our pipeline as described in Materials and methods.

**Fig 1.**
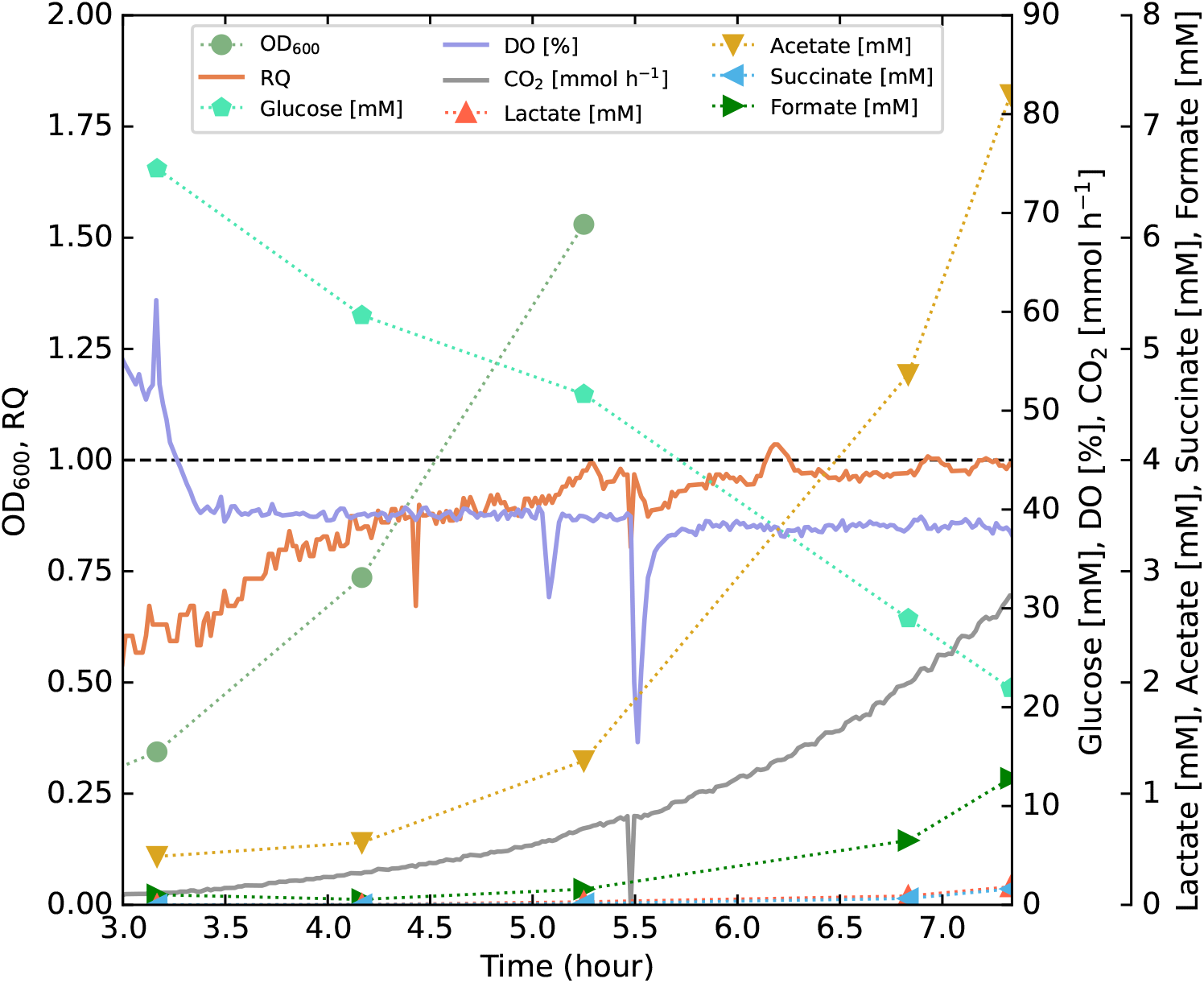
Time-course fermentation profile. Fermentation profile of *E. coli* K-12 MG1655 growing in a minimal glucose medium showing the OD_600_ [unitless], glucose, formate, acetate, and succinate concentrations [mM], as well as CO_2_ in the off-gas [mmol h^−1^], dissolved oxygen (DO) [%] and respiratory quotient (RQ) [unitless]. Unity is highlighted on the same axis as the RQ for reference. Sampling occurred at the final time point (∼7.5 h). Data used for plotting can be found in S4 File. The same figure, except plotted from *t* = 0, can be seen in S4 Fig.

### Macromolecular biomass composition

The measured biomass composition is presented in Table 1, along with the gold-standard reference values by Neidhardt and the biomass distribution reported by Beck *et al*. [15, 38]. While the data from Beck *et al*. are biomass measurements of *E. coli* K-12 MG1655 grown in comparable conditions, the data by Neidhardt contains the biomass profile of *E. coli* B/r based on a combination of experimental data and estimated data from a collection of literature sources. Although not from the same strain, these latter data are routinely used to construct the BOF of *E. coli* GEMs [9, 24] and are commonly employed as a benchmark to evaluate the quality and coverage of biomass composition quantification [15, 19].

**Table 1.** Macromolecular biomass composition. Overview of the average mass fractions [g gCDW^−1^] for the experimentally measured biomass components of *E. coli* K-12 MG1655. Also presented are the previously published *E. coli* biomass compositions by Neidhardt [38] and Beck *et al*. [15]. Standard deviations are included in parentheses. Abbreviations: NA for not applicable, n.d. for not detected, - for not measured/reported, KDO for 2-keto-3-deoxy-octonate.

| Component | This work | Neidhardt | Beck et al. |
| --- | --- | --- | --- |
| **Protein** | **0.54 (0.02)** | **0.550** | **0.35 (0.01)** |
| Alanine | 0.046 (0.004) | 0.039 | 0.025 (0.001) |
| Arginine | 0.052 (0.006) | 0.044 | 0.024 (0.001) |
| Asparagine/Aspartate | 0.052 (0.009) | 0.052 | 0.035 (0.001) |
| Cysteine | 0.010 (NA) | 0.009 | 0.004 (NA) |
| Glutamine/Glutamate | 0.049 (0.007) | 0.068 | 0.049 (0.002) |
| Glycine | 0.025 (0.003) | 0.033 | 0.023 (0.001) |
| Histidine | 0.012 (0.003) | 0.012 | 0.0070 (0.0003) |
| Isoleucine | 0.026 (0.002) | 0.031 | 0.017 (0.001) |
| Leucine | 0.050 (0.007) | 0.048 | 0.026 (0.001) |
| Lysine | 0.036 (0.002) | 0.042 | 0.024 (0.001) |
| Methionine | 0.018 (NA) | 0.019 | 0.011 (NA) |
| Phenylalanine | 0.028 (0.002) | 0.026 | 0.018 (0.002) |
| Proline | 0.022 (NA) | 0.020 | 0.013 (0.001) |
| Serine | 0.021 (0.004) | 0.018 | 0.014 (0.002) |
| Threonine | 0.028 (0.001) | 0.024 | 0.0172 (0.0004) |
| Tryptophan | 0.012 (NA) | 0.010 | 0.008 (NA) |
| Tyrosine | 0.016 (0.003) | 0.021 | 0.014 (0.003) |
| Valine | 0.033 (0.003) | 0.040 | 0.022 (0.001) |
| **RNA** | **0.190 (0.004)** | **0.205** | **0.172 (0.005)** |
| AMP | 0.050 (NA) | 0.054 | 0.045 (NA) |
| GMP | 0.064 (NA) | 0.070 | 0.058 (NA) |
| CMP | 0.040 (NA) | 0.038 | 0.036 (NA) |
| UMP | 0.037 (NA) | 0.042 | 0.033 (NA) |
| **DNA** | **0.013 (0.003)** | **0.031** | **0.010 (0.001)** |
| dAMP | 0.003 (NA) | 0.008 | 0.002 (NA) |
| dGMP | 0.004 (NA) | 0.008 | 0.003 (NA) |
| dCMP | 0.003 (NA) | 0.007 | 0.002 (NA) |
| dTMP | 0.003 (NA) | 0.007 | 0.002 (NA) |
| **Carbohydrate** | **0.0312 (0.0005)** | **0.057** | **0.042 (0.002)** |
| Glucose | 0.0224 (0.0002) | 0.028 | - |
| Glucosamine | 0.0053 (0.0002) | 0.003 | - |
| Galactose | 0.0036 (0.0004) | - | - |
| Rhamnose | n.d. | 0.001 | - |
| Heptose | - | 0.005 | - |
| KDO | - | 0.006 | - |
| N-acetylglucosamine | n.d. | 0.006 | - |
| N-acetylmuramic acid | - | 0.008 | - |
| **Lipid** | **0.061 (0.002)** | **0.091** | **0.067 (0.006)** |
| **Others** | - | 0.066 | - |
| *Sum* | **0.835** | **1.000** | **0.643** |

Constituting the bulk of overall biomass, both the levels of protein (54%) and RNA (19.0%) were found to be closely comparable to those of Neidhardt (at 55.0% and 20.5%, respectively). The relative distribution of individual ribonucleotides was also close to Neidhardt, although we observe higher levels of CMP, similar to what Beck *et al*. reported. The quantities of the majority of individual amino acids agreed well with the estimated profile by Neidhardt, whereas the amounts of glutamate/glutamine, glycine, and tyrosine were found to be noticeably lower. Both the levels of lipids (6.1%) and DNA (1.3%) were measured to be slightly lower than the values reported by Neidhardt, a similar finding to that of Beck *et al*. [15]. The contents of the carbohydrate monomer glucose agreed well with that of Neidhardt, whereas the glucosamine amount was found to be significantly higher. We quantified minor amounts of galactose (0.36%) not reported by Neidhardt, in agreement with well-characterized strain-dependent differences in the composition of the outer core region of LPS in *E. coli* [48, 49]. The absence of any detected rhamnose could also be attributed to these strain-specific variations, or simply that the levels were below the detection limit.

### Changes in biomass objective function stoichiometries affects the genome-scale metabolism

We initiated the construction of the eBOF by scaling the stoichiometric coefficients of the biomass precursors in *i*ML1515 to reflect our measurements of the biomass composition. Following this, the resulting biomass now accounts for 91.6% of the total cellular dry mass, a marked improvement on recent work on *E. coli* [15, 19]. We combined this subset of quantified biomass components with the compounds from the *i*ML1515 mBOF that were not measured in this pipeline (i.e. inorganic ions, metabolites, cofactors, and coenzymes), normalizing to obtain a molar mass of 1 g mmol^−1^.

To assess the change in stoichiometric coefficients of eBOF, we calculated their relative change from mBOF (*S*_*r*_), as defined in Eq. 2. While many biomass coefficients were left unmodified (*S*_*r*_ ≈ 1.0), a considerable proportion of biomass components have significantly altered their relative amounts in eBOF (Fig 2A). In fact, we find that ∼ 14.8 % (w/w) of biomass was reallocated from mBOF to eBOF (see S1 Fig for details). These changes largely coincide with the differences from the biomass measurements of Neidhardt (Table 1). The contents of deoxyribonucleotides, certain amino acids (glycine, glutamate, and glutamine), and lipid components displays the largest decrease, while the peptidoglycan and LPS precursor amounts show the greatest increase.

**Fig 2.**
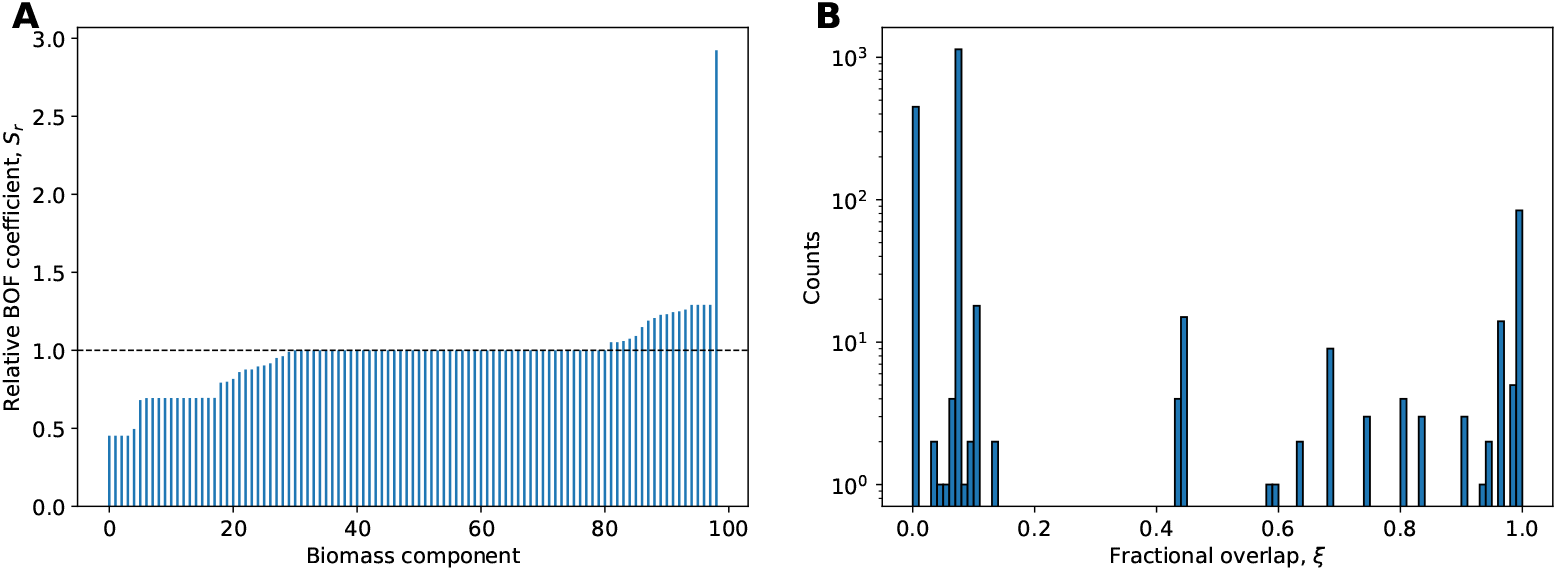
Adjustments of BOF stoichiometries impact attainable flux ranges. (A) Rank-ordered, relative value of BOF coefficients, *S*_*r*_ (Eq. 2), of eBOF compared to mBOF. (B) Histogram of fractional overlap of attainable flux ranges, *ξ* (Eq. 1), for all model reactions using mBOF versus eBOF.

We compared the minimal and maximal reaction fluxes (FVA, 100% optimality constraint), with mBOF and then eBOF as an objective, to give an unbiased overview of the consequences of change in BOF stoichiometries. For every reaction, we calculated the fractional overlap in flux range (*ξ*) as defined in Eq. 1, the distribution of which is presented in Fig 2B. We see that a significant proportion of reactions have no overlap in flux ranges between the two BOFs (*ξ* ≈ 0.0). The remaining reactions are largely clustered in two groupings; one where only half of the flux ranges are overlapping (*ξ* ≈ 0.5), and another where the feasible fluxes are comparable and overlapping (*ξ* ≈ 1.0). The non-overlapping flux ranges overall reflect the reallocation of biomass precursors from mBOF to eBOF. We see a near threefold increase in attainable fluxes of the reactions necessary for the biosynthesis of LPS. For instance, the reaction MCOATA (malonyl-CoA-ACP transacylase) required for the production of myristic acid of the lipid A component of LPS, shifted its flux range from 0.0287 − 0.0289 to 0.0830 − 0.0830 mmol gCDW^−1^ h^−1^ (Table 2). Similarly, the elevated levels of peptidoglycan precursors resulted in a proportional increase in the corresponding biosynthetic flux (e.g. reaction UAGDP, UDP-N-acetylglucosamine diphosphorylase). On the other hand, the reduction in DNA content significantly shifted the flux range of multiple reactions in the nucleotide precursor metabolism (Table 2). The latter can be exemplified by a stark decline in feasible flux of TMDS (thymidylate synthase) from 0.0218 − 0.0218 to 0.0098 − 0.0098 mmol gCDW^−1^ h^−1^ necessary for the biosynthesis of dTMP, as well as a corresponding change in flux range for pyruvate synthase from 0.0938 − 0.0946 to 0.0440 − 0.0441 mmol gCDW^−1^ h^−1^ required for the regeneration of reduced flavodoxin used for converting ribonucleotides into deoxyribonucleotides (Table 2).

**Table 2.** Significant changes in attainable flux ranges. Examples of reactions in *i*ML1515 with a significant shift in feasible flux range when using eBOF. The flux ranges for mBOF is also presented, as well as the relative change in center point (CP_*r*_) of the corresponding flux ranges. Abbreviations: MCOATA for malonyl-CoA-ACP transacylase, UAGDP for UDP-N-acetylglucosamine diphosphorylase, POR5 for pyruvate synthase, TMDS for thymidylate synthase.

| Reaction | Pathway | mBOF | eBOF | $CP_r$ |
| --- | --- | --- | --- | --- |
| MCOATA | Membrane Lipid Metabolism | $0.0287 - 0.0289$ | $0.0830 - 0.0830$ | 2.88 |
| UAGDP | Cell Envelope Biosynthesis | $0.0628 - 0.0629$ | $0.1020 - 0.1020$ | 1.62 |
| POR5 | Pyruvate Metabolism | $0.0938 - 0.0946$ | $0.0440 - 0.0441$ | 0.47 |
| TMDS | Nucleotide Salvage Pathway | $0.0218 - 0.0218$ | $0.0098 - 0.0098$ | 0.45 |

## Discussion

The BOF is a central pseudo-reaction in constraint-based metabolic models containing the metabolic precursors required for cellular replication and growth [9, 15, 20, 21, 50, 51]. Currently, most *E. coli* BOFs can trace their lineage back to data compiled by Neidhardt in 1990 [38]; it should be noted that these data are not for a single *E. coli* strain, and are partly based on measurements from a range of literature sources. The biomass composition of an organism, however, can be highly dynamic and dependent on the strain, growth conditions, growth rate, and growth phase [11, 19]. Consequently, exact and reliable quantification is key in order for the model to accurately predict metabolic phenotypes. The pipeline presented here for the quantification of biomass composition addresses this issue.

Using samples of *E. coli* K-12 MG1655 grown in a defined minimal glucose medium, we obtained a total mass coverage of 91.6% under experimental conditions comparable to those of Neidhardt [38]. This is a marked improvement on the mass recovery recently reported by Beck *et al*. [15] of 64.3%, simultaneously enhancing the molecular resolution. Improving the overall coverage not only provides a more accurate picture of the presence and distribution of biomass components, but also alleviates the unfavourable effects of loss-adjustment by normalization needed to assemble a functional BOF. This normalization step is necessary as the BOF quantitatively represents the conversion of biomass precursors in mmol to gCDW of biomass, thus the molecular mass of the biomass must be scaled to 1 g mmol^−1^ to allow for prediction of specific growth rates [10]. Extensive loss-adjustment overestimates the relative proportion of biomass components with a higher recovery. This is particularly evident for protocols with initial extraction steps (e.g. for DNA), in contrast to those without (e.g. for carbohydrates). Aiming at maximizing the mass coverage is therefore critical in order to obtain a biomass composition of satisfactory quality for constraint-based metabolic modeling.

We obtain an overall comparable amino acid profile to that reported by Neidhardt, although a few amino acids (e.g. glutamate/glutamine, glycine, tyrosine) were found to deviate significantly (Table 1). The under-reported quantities of glutamate/glutamine is presumably caused by their conversion to 5-oxoproline at prolonged exposure to high temperatures [52], which is not measured in this experiment. The reason for the observed inconsistency in the levels of glycine and tyrosine is less clear, although Long and Antoniewicz [19] similarly reported noticeably lower levels of glycine relative to Neidhardt.

Several experimental adjustments are available to correct for or avoid these losses. For instance, introducing correction factors by non-linear regression of serial hydrolysis could aid in the quantification of amino acids that vary in their ease of peptide bond cleavage or chemical resistance to acidic thermohydrolysis [53, 54]. Including the detection and measurement of 5-oxoproline could also assist in recovering a more realistic estimate of glutamate/glutamine levels [52]. Alternatively, the use of other acids and shorter hydrolysis times at higher temperatures has been shown to produce good quantitative yields and with limited side effects in the undesirable degradation of amino acids [55, 56].

Although quantified by our HPLC analysis, methionine was only detected in low concentrations, with a lot of variation across repeated measurements. We believe this to be caused by the propensity for thiol-containing amino acids to undergo oxidative deterioration during acidic hydrolysis [57]. This is further substantiated by our inability to detect cysteine, causing us to estimate the contents of both amino acids based on a linear regression of measured amino acid mass fractions and relative amino acid levels of protein-coding genes. Ensuring complete oxidation of methionine and cysteine by the addition of methanesulfonic acid or performic acid would allow for reliable quantification of the more chemically stable products methionine sulphone and cysteic acid [20, 57].

The cellular amounts of DNA should be a fairly stable quantity at higher growth rates [58]. The low levels might therefore hint at inadequate extraction during the DNA quantification. While this can be counteracted by spiking the samples with DNA standards or isotopically labeled DNA to account and correct for procedural sample loss, there remains a high risk of unaccounted-for matrix effects due to the complexity and heterogeneity of biomass. Analytical approaches which do not require initial extraction steps could therefore be considered promising alternatives for quantification of cellular DNA. For instance, Huang *et al*. proposed a method by which the nucleobase levels of complex cellular material are measured using HPLC following vapor and liquid phase hydrolysis to estimate the overall contents of DNA and RNA [59]. As well as providing a unified platform for the simultaneous quantification of both macromolecules, this approach has the added benefit of directly measuring the levels of individual nucleobases, avoiding the need for biased estimates of their relative distribution from the genome.

No N-acetylglucosamine was detected during our measurements, a carbohydrate that is expected to comprise a significant proportion of the peptidoglycan cell wall of prokaryotes [60]. We hypothesize that this is caused by extensive de-N-acetylation of N-acetylglucosamine to glucosamine during the preliminary hydrolysis step [61, 62]. This is further indicated by our measurements of excessive amounts of glucosamine. While it is present in other *Enterobacteriaceae* strains, it is not supposed to be present in the oligosaccharide core of LPS in strain K-12 [63]. The detection of galactose is also in direct accordance with the same strain-specific characteristics, where a single protruding galactose side chain is present in the outer region of the K-12 core type [63]. When constructing eBOF for *i*ML1515, we therefore assumed all detected glucosamine to originate from N-acetylglucosamine, and treated the contents of N-acetylglucosamine and galactose as proxies for peptidoglycan and LPS levels, respectively. Knowing the monomeric stoichiometry of these complex biopolymers, this allowed for seamless integration with the existing BOF of *i*ML1515 by scaling their stoichiometric coefficients based on our experimental measurements (for calculation details, see Supplementary materials S2 File). This approach has the added benefit of implicitly quantifying N-acetylmuramic acid in peptidoglycan, and 2-keto-3-deoxy-octonate (KDO) and heptose in LPS, which were not directly measured in our analysis. Accounting for these additional contributions, we end up with an overall carbohydrate content of 7.3%. Our approach therefore entails an improvement in both coverage and molecular resolution, the latter of which is commonly lacking in carbohydrate quantification for the analysis of biomass compositions [15, 19, 20]. Whereas other work assumed all carbohydrates to be glucose (thus overestimating its contribution) when implementing a BOF based on their experimental measurements [15], our analysis allows for more precise and nuanced differentiation of carbohydrates into individually quantified sugar compounds.

In addition to the carbohydrates listed in Table 1 we measured 5.1% ribose. While this quantity in theory could be employed to verify the levels of RNA [19], the poor stability of ribose at higher temperatures, particularly in strongly acidic conditions [64], hinders its use as a reliable estimate of cellular RNA. The susceptibility of particular carbohydrate monomers to acid-catalyzed thermohydrolysis is therefore an issue that needs to be addressed in future renditions of the pipeline. Whether this can best be counteracted by changing the acid (type and concentration), by varying physical parameters (e.g. time, temperature), by shielding using chemical modifications, or by quantification of degradation products, remains to be determined.

Cells contain a plethora of chemically diverse compounds and biopolymers that are distinct from the major biomass classes measured in this study. For instance, many genera of both Gram-positive and Gram-negative bacteria have been found to synthesize and accumulate protein- or phospholipid-encapsulated intracellular inclusions of polyhydroxyalkanoates (PHAs) [65]. The biomass contents of PHAs vary strongly with environmental conditions such as nitrogen availability [65], and some *Pseudomonas* strains may even contain up to 67% PHA when engineered for its production [66]. Many BOFs also contain metabolites not covered by the major biomass components, such as soluble metabolites, energy equivalents (e.g. ATP), reducing power (e.g. NAD(P)H), and other coenzymes [24]. While their contribution to the overall biomass is minor, often not reaching levels above 5% [51], they are key factors necessary for cellular proliferation and growth. Not only does this group of intracellular metabolites adhere to organism and cell line-specific distributions [67], it also dynamically adjusts itself in response to changing growth and environmental conditions [68]. The integration of these molecules should therefore greatly enhance the quality of GEMs, as well as aid in the discovery of differences in the metabolic phenotype due to alterations in the intracellular pool of soluble metabolites. It is therefore necessary to broaden the range of detected compounds in our pipeline to account for such organism-, condition- and strain-specific characteristics, while simultaneously maintaining high generality to enable the capture of subtle differences in the macro- and micromolecular biomass compositions of physiologically diverse microorganisms.

After adjusting the coefficients according to the experimental measurements, we arrived at a final mass redistribution of ∼ 14.8%. This suggests that the current BOF for *E. coli*, based on older measurements and adapted throughout the years, is well suited for simulating exponential aerobic growth on glucose minimal medium. While interesting in their own right, the changes in BOF coefficients provide minimal information on the effects of the resulting predictions of genome-scale metabolic fluxes. By being the penultimate end-point of biochemical transformations in GEMs, alterations in the stoichiometric coefficients of these biomass precursors should propagate throughout the metabolic network and affect the attainable fluxes of the model reactions. We therefore performed FVA for mBOF and eBOF using the default *i*ML1515 uptake parameters to look at the consequences for the metabolism of this redistribution in mass. We observe that the altered BOF stoichiometries considerably impact the range of feasible fluxes in the model. This is evident from the fractional overlap of reaction flux ranges *ξ* (Fig 2), as well as the relative change in center point of flux ranges (Supplementary Figure S3 Fig). The latter shows that ∼ 46% of the high flux-carrying reactions have changed their center point by more than 10% when using eBOF compared to mBOF. As the biomass compositions of mBOF and eBOF originate from similar experimental conditions, this difference is rather considerable. Additionally, as biomass compositions significantly vary with growth conditions [12], one would expect the impact on model predictions to be even greater when simulating the metabolic phenotype in different environments. This has profound implications for the application of GEMs and emphasizes the importance of condition-specific biomass measurements when attempting to model a particular scenario.

## Conclusion

A detailed and condition-specific BOF is a key element in making accurate predictions of metabolic phenotypes using GEMs. This necessitates high-quality quantification of the biomass composition of the organism in question, and for the experimental condition being modelled. Here, we present a comprehensive analytical pipeline for absolute quantification of the macromolecular biomass composition of *E. coli* K-12 MG1655 for the construction of strain- and condition-specific BOFs. While rather simple and chiefly relying on well-established protocols for measuring the individual macromolecular classes, we achieved marked improvements in coverage and resolution compared to recently published pipelines. The resulting BOF is made available in the GEM *i*ML1515a.

We applied the experimental pipeline to generate eBOF. The comparison with mBOF revealed a largely similar metabolic phenotype for key attributes like growth and uptake rates, yet the FVA showed a shift in the feasible range of many high-throughput reaction fluxes. Our results therefore highlight the importance of the exact formulation of the BOF, and the need for exact experimental determination for more accurate predictions, even for well-studied organisms such as *E. coli* under the most standard of conditions. For less-studied organisms, and under more esoteric conditions, one would reasonably expect the impact of a specifically determined BOF to be dramatically higher.

With this work, we address what we regard to be one of the more pressing subjects in the constraint-based metabolic modeling community: the unfortunate tradition of not allocating resources into experimental determination of biomass composition. We have shown that while it remains time-consuming, it is indeed both important and possible to make these measurements. Additionally, such a biomass determination pipeline opens the possibility to generate multiple biomass compositions under different growth conditions for a given organism. It is our hope that the presented protocols and techniques will be further adapted and improved, and that the measurement of biomass composition will become routine.

## Acknowledgments

The authors would like to thank Siri Stavrum for assisting with the HPLC analysis. Additionally, we would like to thank Kåre Andre Kristiansen, Zdenka Bartosova, and Kanhaiya Kumar for helpful input throughout the project.

## Supporting information

**S1 Fig.**
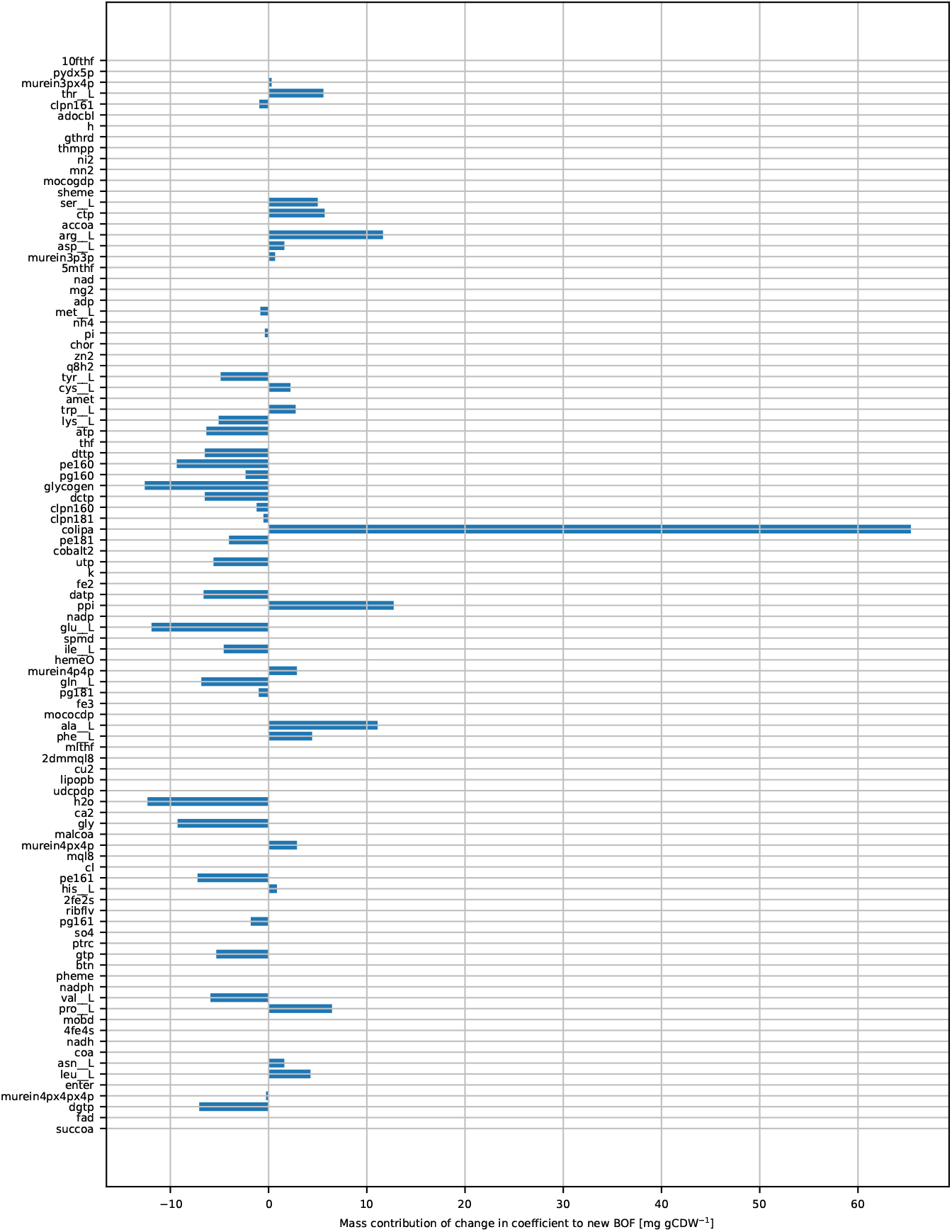
Similarity of the BOFs. The difference between the *i*ML1515 WT BOF (mBOF) and the BOF determined in the manuscript (eBOF), in terms of the mass redistribution [mg g^−1^] resulting from the change in each biomass coefficient. Negative values indicate that the relative amount of the given component was reduced in eBOF compared to mBOF, while positive values indicate that the relative amount of the given component was increased.

**S2 Fig.**
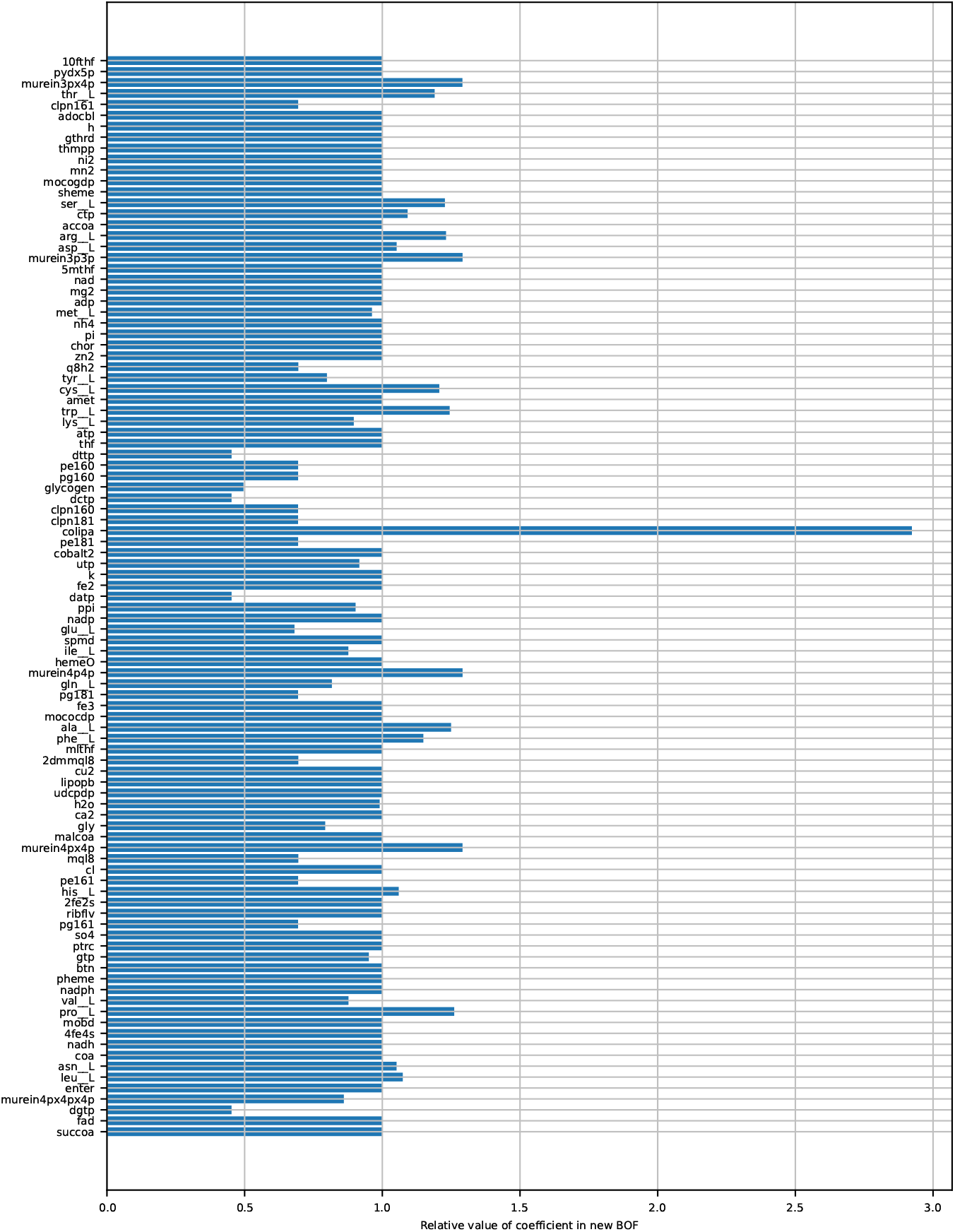
Relative similarity of the BOFs. The difference between mBOF and eBOF in terms of the ratio of each biomass coefficient. Values smaller than unity indicate that the relative amount of the given component was reduced in eBOF compared to mBOF, while values greater than unity indicate that the relative amount of the given component was increased.

**S3 Fig.**
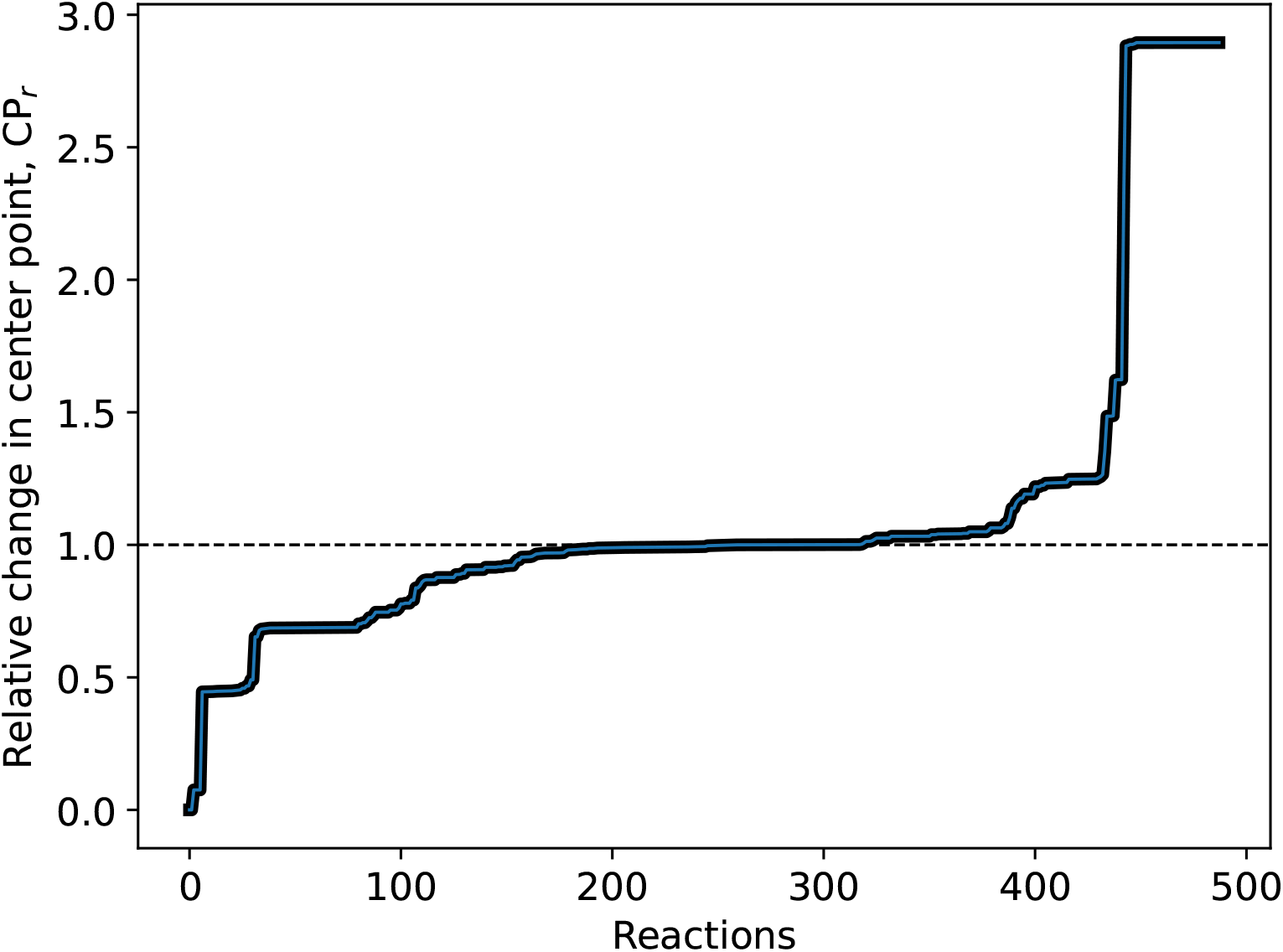
Relative change in center point (CP_*r*_). Relative change in center point (CP_*r*_) of the flux ranges in the high-flux carrying reactions (absolute flux *>* 0.001 mmol gCDW^−1^ h^−1^) when using eBOF versus mBOF. The corresponding flux ranges were calculated by performing a flux variability analysis (FVA) for all reactions in *i*ML1515 at optimal growth phenotype using the default uptake parameters of the model.

**S4 Fig.**
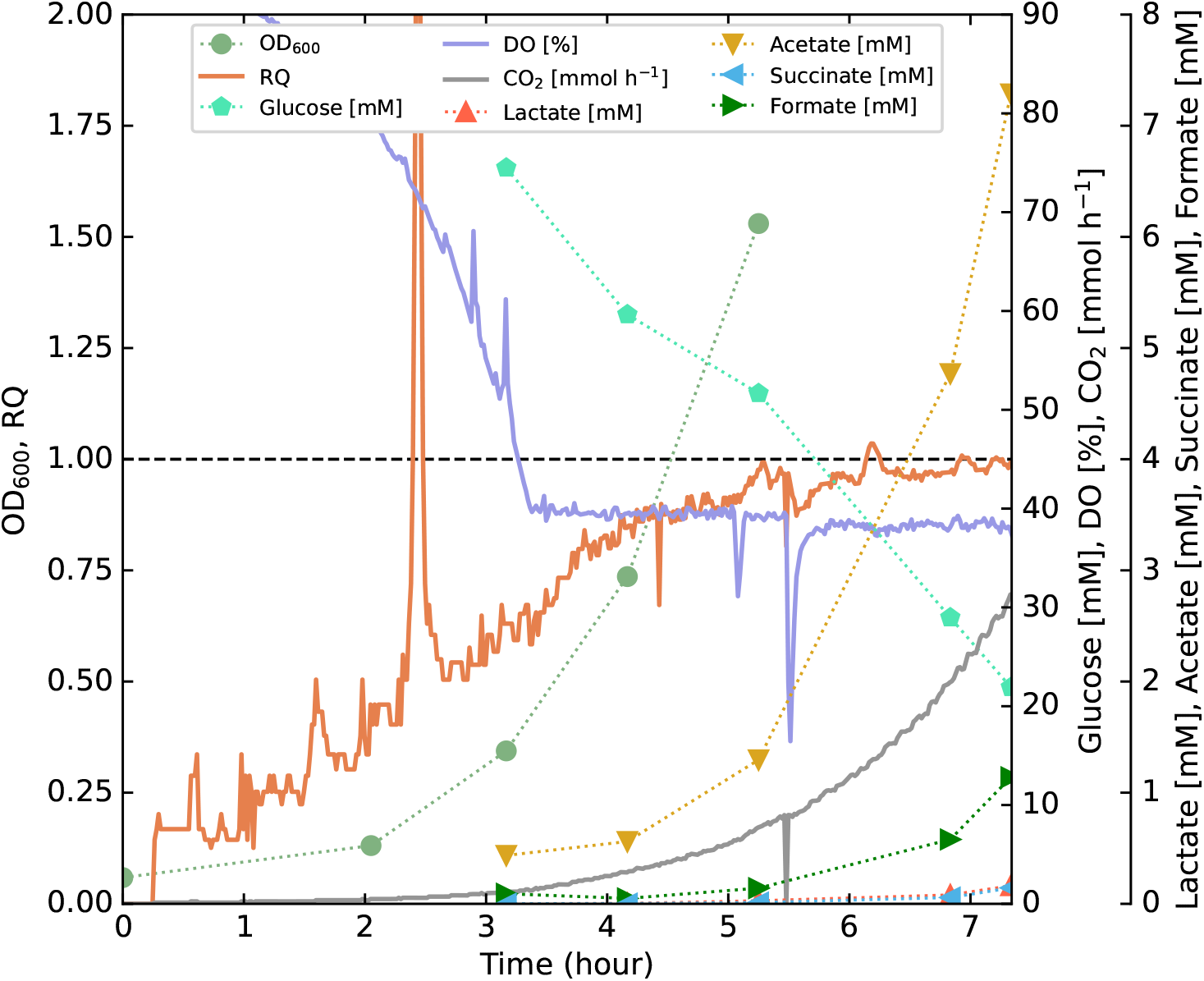
Time-course fermentation profile from time zero. Fermentation profile of *E. coli* K-12 MG1655 growing in a minimal glucose medium showing the OD_600_ [unitless], glucose, formate, acetate, and succinate concentrations [mM], as well as CO_2_ in the off-gas [mmol h^−1^], dissolved oxygen (DO) [%] and respiratory quotient (RQ) [unitless]. Unity is highlighted on the same axis as the RQ for reference. Data used for plotting can be found in S4 File. This is essentially the same plot as Fig 1, except plotted from *t* = 0.

**S1 File. The genome-scale metabolic model *i*ML1515a**. Revised *i*ML1515 model (named *i*ML1515a) with a corrected reaction GALT1, and with the experimentally determined eBOF included.

**S2 File. Descriptions of eBOF construction**. Here, we provide a detailed description on how eBOF was constructed using our experimental measurements.

**S3 File. Experimental measurements and eBOF calculations**. Experimental measurements of the macromolecular biomass composition, as well as the calculations used to construct eBOF for *i*ML1515a.

**S4 File. Fermentor and sampling data used for generating the plots in 1 and S4 Fig** Here we provide the data for fermentor measurements such as off-gas and DO, as well as sampling data of OD and different compound concentrations in the fermentor.

## References

1. Chuang HY, Hofree M, Ideker T. A Decade of Systems Biology. Annual Review of Cell and Developmental Biology. 2010;26(1):721–744. doi:10.1146/annurev-cellbio-100109-104122.

2. Kitano H. Systems biology: A brief overview. Science. 2002;295(5560):1662–1664. doi:10.1126/science.1069492.

3. Bordbar A, Monk JM, King ZA, Palsson BO. Constraint-based models predict metabolic and associated cellular functions. Nature Reviews Genetics. 2014;15(2):107–120. doi:10.1038/nrg3643.

4. Price ND, Reed JL, Palsson B. Genome-scale models of microbial cells: Evaluating the consequences of constraints. Nature Reviews Microbiology. 2004;2(11):886–897. doi:10.1038/nrmicro1023.

5. Varma A, Palsson BO. Metabolic flux balancing: Basic concepts, scientific and practical use. Bio/Technology. 1994;12(10):994–998. doi:10.1038/nbt1094-994.

6. Oberhardt MA, Palsson BØ, Papin JA. Applications of genome scale metabolic reconstructions. Molecular Systems Biology. 2009;5(1):320. doi:10.1038/msb.2009.77.

7. Orth JD, Thiele I, Palsson BØO. What is flux balance analysis? Nature Biotechnology. 2010;28(3):245–248. doi:10.1038/nbt.1614.

8. Schuetz R, Kuepfer L, Sauer U. Systematic evaluation of objective functions for predicting intracellular fluxes in Escherichia coli. Molecular Systems Biology. 2007;3(1):119. doi:10.1038/msb4100162.

9. Feist AM, Palsson BO. The biomass objective function. Current Opinion in Microbiology. 2010;13(3):344–349. doi:https://doi.org/10.1016/j.mib.2010.03.003.

10. Chan SHJ, Cai J, Wang L, Simons-Senftle MN, Maranas CD. Standardizing biomass reactions and ensuring complete mass balance in genome-scale metabolic models. Bioinformatics. 2017;33(22):3603–3609. doi:10.1093/bioinformatics/btx453.

11. Dennis PP, Bremer H. Modulation of Chemical Composition and Other Parameters of the Cell at Different Exponential Growth Rates. EcoSal Plus. 2008;3(1). doi:10.1128/ecosal.5.2.3.

12. Schaechter M, MaalOe O, Kjeldgaard NO. Dependency on Medium and Temperature of Cell Size and Chemical Composition during Balanced Growth of Salmonella typhimurium. Journal of General Microbiology. 1958;19(3):592–606. doi:10.1099/00221287-19-3-592.

13. Taymaz-Nikerel H, Borujeni AE, Verheijen PJT, Heijnen JJ, van Gulik WM. Genome-derived minimal metabolic models for Escherichia coli MG1655 with estimated in vivo respiratory ATP stoichiometry. Biotechnology and Bioengineering. 2010;107(2):369–381. doi:10.1002/bit.22802.

14. Simensen V, Voigt A, Almaas E. High quality genome scale metabolic model of Aurantiochytrium sp. T66. Biotechnology and Bioengineering. 2021; p. bit.27726. doi:10.1002/bit.27726.

15. Beck A, Hunt K, Carlson R. Measuring Cellular Biomass Composition for Computational Biology Applications. Processes. 2018;6(5):38. doi:10.3390/pr6050038.

16. Széliová D, Ruckerbauer DE, Galleguillos SN, Petersen LB, Natter K, Hanscho M, et al. What CHO is made of: Variations in the biomass composition of Chinese hamster ovary cell lines. Metabolic Engineering. 2020;61:288–300. doi:10.1016/j.ymben.2020.06.002.

17. Lakshmanan M, Long S, Ang KS, Lewis N, Lee DY. On the impact of biomass composition in constraint-based flux analysis; 2019. Available from: https://doi.org/10.1101/652040.

18. Dikicioglu D, Kırdar B, Oliver SG. Biomass composition: the “elephant in the room” of metabolic modelling. Metabolomics. 2015;11(6):1690–1701. doi:10.1007/s11306-015-0819-2.

19. Long CP, Antoniewicz MR. Quantifying Biomass Composition by Gas Chromatography/Mass Spectrometry. Analytical Chemistry. 2014;86(19):9423–9427. doi:10.1021/ac502734e.

20. Széliová D, Schoeny H, Knez Š, Troyer C, Coman C, Rampler E, et al. Robust Analytical Methods for the Accurate Quantification of the Total Biomass Composition of Mammalian Cells. In: Deepak Nagrath, editor. Methods in Molecular Biology. vol. 2088. Ann Arbor, MI, USA: Humana Press; 2020. p. 119–160. Available from: http://link.springer.com/10.1007/978-1-0716-0159-4{_}7.

21. Schulz C, Kumelj T, Karlsen E, Almaas E. Genome-scale metabolic modelling when changes in environmental conditions affect biomass composition. PLoS Computational Biology. 2021;17(5):e1008528. doi:10.1371/journal.pcbi.1008528.

22. Bartosova Z, Gonzalez SV, Voigt A, Bruheim P. High Throughput Semiquantitative UHPSFC–MS/MS Lipid Profiling and Lipid Class Determination. Journal of Chromatographic Science. 2021;2021:1–11. doi:10.1093/chromsci/bmaa121.

23. Rühmann B, Schmid J, Sieber V. Fast carbohydrate analysis via liquid chromatography coupled with ultra violet and electrospray ionization ion trap detection in 96-well format. Journal of Chromatography A. 2014;1350:44–50. doi:10.1016/j.chroma.2014.05.014.

24. Monk JM, Lloyd CJ, Brunk E, Mih N, Sastry A, King Z, et al. iML1515, a knowledgebase that computes Escherichia coli traits. Nature Biotechnology. 2017;35(10):904–908. doi:10.1038/nbt.3956.

25. Hong RS, Hwang KH, Kim S, Cho HE, Lee HJ, Hong JT, et al. Survey of ERETIC2 NMR for quantification. Journal of the Korean Magnetic Resonance Society. 2013;17(2):98–104. doi:10.6564/JKMRS.2013.17.2.098.

26. Søgaard CK, Blindheim A, Røst LM, Petrović V, Nepal A, Bachke S, et al. “Two hits - one stone”; increased efficacy of cisplatin-based therapies by targeting PCNA’s role in both DNA repair and cellular signaling. Oncotarget. 2018;9(65):32448–32465. doi:10.18632/oncotarget.25963.

27. Isbell HS, Ward Pigman. THE OXIDATION OF ALPHA AND BETA GLUCOSE AND A STUDY OF THE ISOMERIC FORMS OF THE SUGAR IN SOLUTION. B S Jour Research. 1932;8((RP418)):327.

28. Wishart DS, Knox C, Guo AC, Eisner R, Young N, Gautam B, et al. HMDB: a knowledgebase for the human metabolome. Nucleic Acids Research. 2009;37(Database):D603–D610. doi:10.1093/nar/gkn810.

29. Wishart DS, Jewison T, Guo AC, Wilson M, Knox C, Liu Y, et al. HMDB 3.0—The Human Metabolome Database in 2013. Nucleic Acids Research. 2012;41(D1):D801–D807. doi:10.1093/nar/gks1065.

30. Wishart DS, Feunang YD, Marcu A, Guo AC, Liang K, Vázquez-Fresno R, et al. HMDB 4.0: the human metabolome database for 2018. Nucleic Acids Research. 2018;46(D1):D608–D617. doi:10.1093/nar/gkx1089.

31. Chenomx INC. Chenomx NMR Mixture analysis; 2021. Available from: https://www.chenomx.com/.

32. Fan TWM. Metabolite profiling by one-and two-dimensional NMR analysis of complex mixtures. Progress in Nuclear Magnetic Resonance Spectroscopy. 1996;28(2):161–219. doi:10.1016/0079-6565(95)01017-3.

33. Noble JE, Knight AE, Reason AJ, Di Matola A, Bailey MJA. A Comparison of Protein Quantitation Assays for Biopharmaceutical Applications. Molecular Biotechnology. 2007;37(2):99–111. doi:10.1007/s12033-007-0038-9.

34. Holt L, Milligan B, Roxburgh C. Aspartic Acid, Asparagine, Glutamic Acid, and Glutamine Contents of Wool and two Derived Protein Fractions. Australian Journal of Biological Sciences. 1971;24(3):509. doi:10.1071/BI9710509.

35. Benthin S, Nielsen J, Villadsen J. A simple and reliable method for the determination of cellular RNA content. Biotechnology Techniques. 1991;5(1):39–42. doi:10.1007/BF00152753.

36. Desjardins P, Conklin D. NanoDrop Microvolume Quantitation of Nucleic Acids. Journal of Visualized Experiments. 2010;45(-1):1–5. doi:10.3791/2565.

37. Clark K, Karsch-Mizrachi I, Lipman DJ, Ostell J, Sayers EW. GenBank. Nucleic acids research. 2016;44(D1):D67–D72. doi:10.1093/nar/gkv1276.

38. Neidhardt FC, Ingraham JL, Schaechter M. Physiology of the Bacterial Cell: A Molecular Approach. 20th ed. MA: Sinauer Associates Sunderland, MA; 1990.

39. Wright MH, Adelskov J, Greene AC. Bacterial DNA Extraction Using Individual Enzymes and Phenol/Chloroform Separation †. Journal of Microbiology & Biology Education. 2017;18(2):1–3. doi:10.1128/jmbe.v18i2.1348.

40. Rühmann B, Schmid J, Sieber V. Automated modular high throughput exopolysaccharide screening platform coupled with highly sensitive carbohydrate fingerprint analysis. Journal of Visualized Experiments. 2016;2016(110):1–14. doi:10.3791/53249.

41. Bligh EG, Dyer WJ. A rapid method of total lipid extraction and purification. Canadian journal of biochemistry and physiology. 1959;37(8):911–917. doi:10.1139/o59-099.

42. Skedsmo FS, Malachin G, Våge DI, Hammervold MM, Salvesen Ø, Ersdal C, et al. Demyelinating polyneuropathy in goats lacking prion protein. The FASEB Journal. 2020;34(2):2359–2375. doi:10.1096/fj.201902588R.

43. Burgard AP, Vaidyaraman S, Maranas CD. Minimal reaction sets for Escherichia coli metabolism under different growth requirements and uptake environments. Biotechnology Progress. 2001;17(5):791–797. doi:10.1021/bp0100880.

44. Mahadevan R, Schilling CH. The effects of alternate optimal solutions in constraint-based genome-scale metabolic models. Metabolic Engineering. 2003;5(4):264–276. doi:10.1016/j.ymben.2003.09.002.

45. MathWorks. Matlab 2020a; 2020. Available from: https://www.mathworks.com/.

46. Heirendt L, Arreckx S, Pfau T, Mendoza SN, Richelle A, Heinken A, et al. Creation and analysis of biochemical constraint-based models using the COBRA Toolbox v.3.0. Nature Protocols. 2019;14(3):639–702. doi:10.1038/s41596-018-0098-2.

47. Cplex IBMI. V12. 1: User’s Manual for CPLEX. International Business Machines Corporation. 2009;46(53):157.

48. Washizaki A, Yonesaki T, Otsuka Y. Characterization of the interactions between *Escherichia coli* receptors, LPS and OmpC, and bacteriophage T4 long tail fibers. MicrobiologyOpen. 2016;5(6):1003–1015. doi:10.1002/mbo3.384.

49. Raetz CRH, Whitfield C. Lipopolysaccharide endotoxins; 2002. Available from: /pmc/articles/PMC2569852//pmc/articles/PMC2569852/?report= abstracthttps://www.ncbi.nlm.nih.gov/pmc/articles/PMC2569852/.

50. Folsom JP, Carlson RP. Physiological, biomass elemental composition and proteomic analyses of Escherichia coli ammonium-limited chemostat growth, and comparison with iron-and glucose-limited chemostat growth. Microbiology. 2015;161(8):1659–1670. doi:10.1099/mic.0.000118.

51. Lachance JC, Lloyd CJ, Monk JM, Yang L, Sastry AV, Seif Y, et al. BOFdat: Generating biomass objective functions for genome-scale metabolic models from experimental data. PLOS Computational Biology. 2019;15(4):e1006971. doi:10.1371/journal.pcbi.1006971.

52. Shih FF. Amsterdam-Printed in The Netherlands CHROM; 1985.

53. Darragh AJ, Moughan PJ. The effect of hydrolysis time on amino acid analysis. Journal of AOAC International. 2005;88(3):888–893. doi:10.1093/jaoac/88.3.888.

54. Lapierre H, Binggeli S, Sok M, Pellerin D, Ouellet DR. Estimation of correction factors to determine the true amino acid concentration of protein after a 24-hour hydrolysis. Journal of Dairy Science. 2019;102(2):1205–1212. doi:10.3168/jds.2018-15392.

55. Chiou SH, Wang KT. Simplified protein hydrolysis with methanesulphonic acid at elevated temperature for the complete amino acid analysis of proteins. Journal of Chromatography A. 1988;448(C):404–410. doi:10.1016/S0021-9673(01)84603-3.

56. Marino R, Iammarino M, Santillo A, Muscarella M, Caroprese M, Albenzio M. Technical note: Rapid method for determination of amino acids in milk. Journal of Dairy Science. 2010;93(6):2367–2370. doi:10.3168/jds.2009-3017.

57. Albert C, Sára P, Lóki K, Visi ÉV, Salamon S, Csapó J, et al. Investigation of performic acid oxidation in case of thiol-containing amino acid enantiomers. Acta Agraria Kaposváriensis. 2006;10(2):349–354.

58. Pramanik J, Keasling JD. Stoichiometric model ofEscherichia coli metabolism: Incorporation of growth-rate dependent biomass composition and mechanistic energy requirements. Biotechnology and Bioengineering. 1997;56(4):398–421. doi:10.1002/(SICI)1097-0290(19971120)56:4¡398::AID-BIT6¿3.0.CO;2-J.

59. Huang Q, Kaiser K, Benner R. A simple high performance liquid chromatography method for the measurement of nucleobases and the RNA and DNA content of cellular material. Limnology and Oceanography: Methods. 2012;10(8):608–616. doi:10.4319/lom.2012.10.608.

60. Vollmer W, Blanot D, De Pedro MA. Peptidoglycan structure and architecture. FEMS Microbiology Reviews. 2008;32(2):149–167. doi:10.1111/j.1574-6976.2007.00094.x.

61. Einbu A, Vårum KM. Depolymerization and de-N-acetylation of chitin oligomers in hydrochloric acid. Biomacromolecules. 2007;8(1):309–314. doi:10.1021/bm0608535.

62. Einbu A, Vårum KM. Characterization of chitin and its hydrolysis to GlcNAc and GlcN. Biomacromolecules. 2008;9(7):1870–1875. doi:10.1021/bm8001123.

63. Ebbensgaard A, Mordhorst H, Aarestrup FM, Hansen EB. The role of outer membrane proteins and lipopolysaccharides for the sensitivity of escherichia coli to antimicrobial peptides. Frontiers in Microbiology. 2018;9(SEP):2153. doi:10.3389/fmicb.2018.02153.

64. Larralde R, Robertscn MP, Miller SL. Rates of decomposition of ribose and other sugars: Implications for chemical evolution. Proceedings of the National Academy of Sciences of the United States of America. 1995;92(18):8158–8160. doi:10.1073/pnas.92.18.8158.

65. Raza ZA, Abid S, Banat IM. Polyhydroxyalkanoates: Characteristics, production, recent developments and applications; 2018.

66. Mozejko-Ciesielska J, Szacherska K, Marciniak P. Pseudomonas Species as Producers of Eco-friendly Polyhydroxyalkanoates. Journal of Polymers and the Environment. 2019;27(6):1151–1166. doi:10.1007/s10924-019-01422-1.

67. Røst LM, Thorfinnsdottir LB, Kumar K, Fuchino K, Langørgen IE, Bartosova Z, et al. Absolute quantification of the central carbon metabolome in eight commonly applied prokaryotic and eukaryotic model systems. Metabolites. 2020;10(2):74. doi:10.3390/metabo10020074.

68. Kumar K, Venkatraman V, Bruheim P. Adaptation of central metabolite pools to variations in growth rate and cultivation conditions in Saccharomyces cerevisiae. Microbial Cell Factories. 2021;20(1):1–16. doi:10.1186/s12934-021-01557-8.

