## Supplementary material for "Quantification of macromolecular biomass composition for constraint-based metabolic modeling": S2 File

### Supporting information S2

#### Constructing the biomass objective function

As the BOF quantitatively describes the formation of biomass in gCDW from molar equivalents of biomass precursors, the molecular mass of biomass needs to be normalized to $1\,\mathrm{g\,mmol^{-1}}$. The experimental measurements therefore need to be converted to stoichiometric coefficients in units of $\mathrm{mmol\,gCDW^{-1}}$ (i.e. molar fractions), and appropriately scaled in order for the model to accurately predict specific growth rates [1]. Here, we provide a detailed description on how we constructed eBOF for *i*ML1515a using our experimental measurements. The referenced data and calculations are all made available in S3 File.

##### Protein

The measured amino acid concentrations were first converted to mass fractions, then molar fractions by dividing by their molar masses. As certain amino acids either were partially or close-to fully degraded during this preliminary boiling step, their mass fractions were estimated based on a linear regression of the measured amino acid levels and the relative amino acid distribution in the protein-coding genes of *Escherichia coli* K-12 MG1655.

##### RNA

The concentrations were converted to mass fractions and split between the four ribonucleotides adenosine 5'-monophosphate (AMP), guanosine 5'-monophosphate (GMP), uridine 5'-monophosphate (UMP), and cytidine 5'-monophosphate (CMP) according to the composition of rRNA-encoding genes in *E. coli* strain K-12 MG1655. Each mass fraction was subsequently converted to molar fractions by dividing by their molar masses.

##### DNA

The overall concentration was subdividied into the individual deoxyribonucleotides deoxyadenosine 5'-monophosphate (dAMP), deoxyguanosine 5'-monophosphate (dGMP), thymidine 5'-monophosphate (dTMP), and deoxycytidine 5'-monophosphate (dCMP) in accordance with the relative distribution in the genome sequence of *E. coli* strain K-12 MG1655. These concentrations was further converted to molar fractions by dividing the corresponding mass fractions by their molar masses.

##### Carbohydrate

The measured concentrations were converted to mass fractions while accounting for the additional mass of water from the initial hydrolysis step. These mass fractions were subsequently converted to molar fractions by dividing by their molar masses.

The carbohydrates in mBOF are constituents of three distinct biopolymers: lipopolysaccharide (LPS), peptidoglycan, and glycogen. LPS is represented in *i*ML1515 by a single metabolite: colipa_e, and its content was estimated based on the measured levels of galactose which is present in unimolecular amounts in the outer core of the LPS [2, 3].

We discovered that the model erroneously implemented the responsible galactosyltransferase reaction (GALT1) as it utilized UDP-glucose as a sugar donor, rather than UDP-galactose. Having fixed this, *i*ML1515a is now able to correctly incorporate galactose during the biosynthesis of LPS. Peptidoglycan is represented by five precursor molecules (murein3p3p_p, murein3px4p_p, murein4p4p_p, murein4px4p_p, and murein4px4px4p_p) with varying amounts of the alternating amino sugars N-acetylglucosamine (GlcNAc) and N-acetyl muramic acid (MurNAc), and amino acids L/D-alanine, D-glutamate, and *meso*-diaminopimelate. Due to the absence of glucosamine (GlcN) in the LPS of *E. coli* strain K-12 MG1655 [2, 3], we assumed that all quantified GlcN originated from de-N-acetylated GlcNAc. The relative composition of these five precursor molecules were assumed unchanged and their total amounts were appropriately scaled using our GlcN measurements. Glycogen is a biopolymer of glucose molecules and is represented in *i*ML1515 as a single compound glycogen_c. Correcting for the glucose content of the estimated LPS levels, the remaining glucose is assumed to solely originate from glycogen.

#### Lipid

In *i*ML1515, lipids are represented as phosphatidylethanolamine (PE), phosphatidylglyerol (PG), and cardiolipin (CL). Each of these lipid classes consists of three individual lipid compounds with a specific fatty acid composition. Using the relative distribution in mBOF, we scaled the amounts of each of these compounds to reflect that of our total lipid measurements. We also accounted for the levels of the coenzymes ubiquinol-8 (q8h2_c), menaquinol-8 (mql8_c), and 2-demethylmenaquinol-8 (2dmmql8_c) as their hydrophobicity make them readily soluble in chloroform [4]. While the membrane anchor lipid A is a major component of bacterial LPS [3], its extraction efficiency is highly variable due to the hydrophilic nature of the oligosaccharide core [5]. Consequently, we assumed that negligible amounts of the LPS would be obtained in our lipid extraction procedure and decided not to account for this when scaling the lipid portion of the BOF.

#### Normalizing and constructing eBOF

We decided to include the levels of the biomass components in mBOF that were not directly measured in this work (e.g. inorganic ions, metabolites). Keeping this subset constant, we normalized the levels of the quantified biomass components to obtain a molecular mass of biomass of $1\,\mathrm{g\,mmol^{-1}}$. Both the growth-associated maintenance (GAM) and the non-growth associated maintenance (NGAM) terms were left unchanged.

### References


[1] Chan SHJ, Cai J, Wang L, Simons-Senftle MN, Maranas CD. Standardizing biomass reactions and ensuring complete mass balance in genome-scale metabolic models. Bioinformatics. 2017;33(22):3603–3609. doi:10.1093/bioinformatics/btx453.

[2] Washizaki A, Yonesaki T, Otsuka Y. Characterization of the interactions between <i>Escherichia coli</i> receptors, LPS and OmpC, and bacteriophage T4 long tail fibers. MicrobiologyOpen. 2016;5(6):1003–1015. doi:10.1002/mbo3.384.

[3] Raetz CRH, Whitfield C. Lipopolysaccharide endotoxins; 2002. Available from: /pmc/articles/PMC2569852//pmc/articles/PMC2569852/?report=abstracthttps://www.ncbi.nlm.nih.gov/pmc/articles/PMC2569852/.

[4] Sévin DC, Sauer U. Ubiquinone accumulation improves osmotic-stress tolerance in Escherichia coli. Nature Chemical Biology. 2014;10(4):266–272. doi:10.1038/nchembio.1437.

[5] Furse S, Egmond MR, Killian JA. Isolation of lipids from biological samples; 2015. Available from: https://www.tandfonline.com/action/journalInformation?journalCode=imbc20.
